# Phenotypic variation in two widespread Caribbean corals — *Porites astreoides* and *P. divaricata* — between mangrove and lagoon habitats

**DOI:** 10.1101/2020.06.30.181172

**Authors:** Karina Scavo Lord, Anna Barcala, Hannah E. Aichelman, Nicola G. Kriefall, Chloe Brown, Lauren Knasin, Riley Secor, Cailey Tone, Laura Tsang, John R. Finnerty

## Abstract

As coral reefs experience dramatic declines in coral cover throughout the tropics, there is an urgent need to understand the role that non-reef habitats such as mangroves play in the ecological niche of corals. Mangrove habitats present a challenge to reef-dwelling corals as they can differ dramatically from adjacent reef habitats with respect to key environmental parameters such as temperature and light. As variation in temperature and light within reef habitats is known to drive intraspecific differences in coral phenotype, we hypothesized that coral species which can exploit both reef and mangrove habitats will exhibit predictable differences in phenotype between habitats. To investigate how intraspecific variation, driven by either local adaptation or phenotypic plasticity, might enable particular coral species to exploit these two qualitatively different habitat types, we compared the phenotypes of two widespread Caribbean corals — *Porites divaricata* and *P. astreoides—* in mangrove versus lagoon habitats on Turneffe Atoll, Belize. We document significant differences in colony size, color, structural complexity, and corallite morphology between habitats. In every instance, the difference between mangrove and lagoon corals was consistent in *P. divaricata* and *P. astreoides*. This study is the first to document intraspecific phenotypic diversity in corals occupying mangrove versus patch-reef habitats, and it provides a foundation for understanding why some “reef coral” species can exploit mangroves, while others cannot.

## 1. INTRODUCTION

Globally, coral reefs are declining at an alarming rate due to the effects of climate change, especially rising sea surface temperatures and ocean acidification (Hughes et al. 2017), as well as a myriad of local anthropogenic stressors (Ban et al. 2014). However, reefs are not the only habitat in which corals can live. Many coral species are habitat generalists that can thrive in a multitude of different environments in addition to coral reefs, including mangrove forests (Yates et al. 2014, Hernández Fernández 2015, Rogers 2017, Bengtsson et al. 2019, Camp et al. 2019) and seagrass beds (Camp et al. 2016, Lohr et al. 2017). For example, around half of the approximately 75 coral species that occur on Caribbean reefs also occur in mangrove habitats (Scavo Lord et al. 2020).

The ability to live and reproduce in a range of habitats or under a multitude of conditions could substantially increase species survival rate in this period of rapid environmental change and increased environmental variability. Indeed, in many locations throughout the Caribbean, species that were “competitively dominant” during prior periods of environmental stability are being replaced by “weedy” species that can survive across a range of habitats (Darling et al. 2012, Darling et al. 2013). For example, the iconic Caribbean branching corals, *Acropora cervicornis* and *A. palmata*, were able to grow rapidly and outcompete other corals under the relatively stable conditions that persisted on Caribbean reefs for most of the last few thousand years (Aronson et al. 2002, Darling et al. 2012). However, in recent decades, these Acroporids have suffered severe declines (Aronson & Precht 2001, Precht et al. 2002), while slower-growing “stress tolerant” massive corals (e.g., *Orbicella* spp.) and rapidly-recruiting “weedy corals” (e.g., *Porites* spp.) have increased in prevalence (Edmunds & Carpenter 2001, Gardner et al. 2003, Green et al. 2008). For this reason, the future composition of coral assemblages in the Caribbean and elsewhere may depend, in part, on how increasing environmental variation affects the physiology and fitness of habitat generalists.

One way to investigate how environmental variation affects coral phenotypes is to compare the same coral species across an environmental gradient. For example, in reef habitats, differences in depth (and associated differences in light and flow) have been shown to drive differences in microskeletal anatomy (Ow & Todd 2010, Soto et al. 2018) and color (Gleason 1993). Presumably, this intraspecific variation is advantageous because a generalist that can live in a wide range of depths is at an advantage over a specialist that can only occur in a narrow range of depths.

To investigate how intraspecific variation, driven by either genetic variation or phenotypic plasticity, might enable corals to exploit two qualitatively different habitat types, we compared the phenotypes of two widespread Caribbean corals — *Porites divaricata* and *P. astreoides—* in mangrove versus lagoon habitats on Turneffe Atoll, Belize (Figure 1). These two weedy corals can be found in a wide variety of habitats. The thin finger coral, *P. divaricata*, is common in shallow lee habitats, including the reef crest, seagrass beds, mangrove forests and lagoonal patch reefs (Veron 2000, Hernández Fernández 2015, Camp et al. 2016, Rogers 2017, Bengtsson et al. 2019). This branching coral can vary greatly in overall shape, number and density of branch tips, and color (Figure 2). The mustard hill coral, *P. astreoides*, can be found in all reef habitats at depths between 0-50 m, as well as mangrove forests, seagrass beds, and shallow lagoon patch reefs (Veron 2000, Hernández Fernández 2015, Camp et al. 2016, Rogers 2017, Bengtsson et al. 2019). This species has highly variable morphology and color (Figure 3), existing as a massive, encrusting, or plating form, with a smooth or bumpy surface, and colors ranging from bright yellow to medium gray to dark brown (Veron 2000).

**Figure 1.**
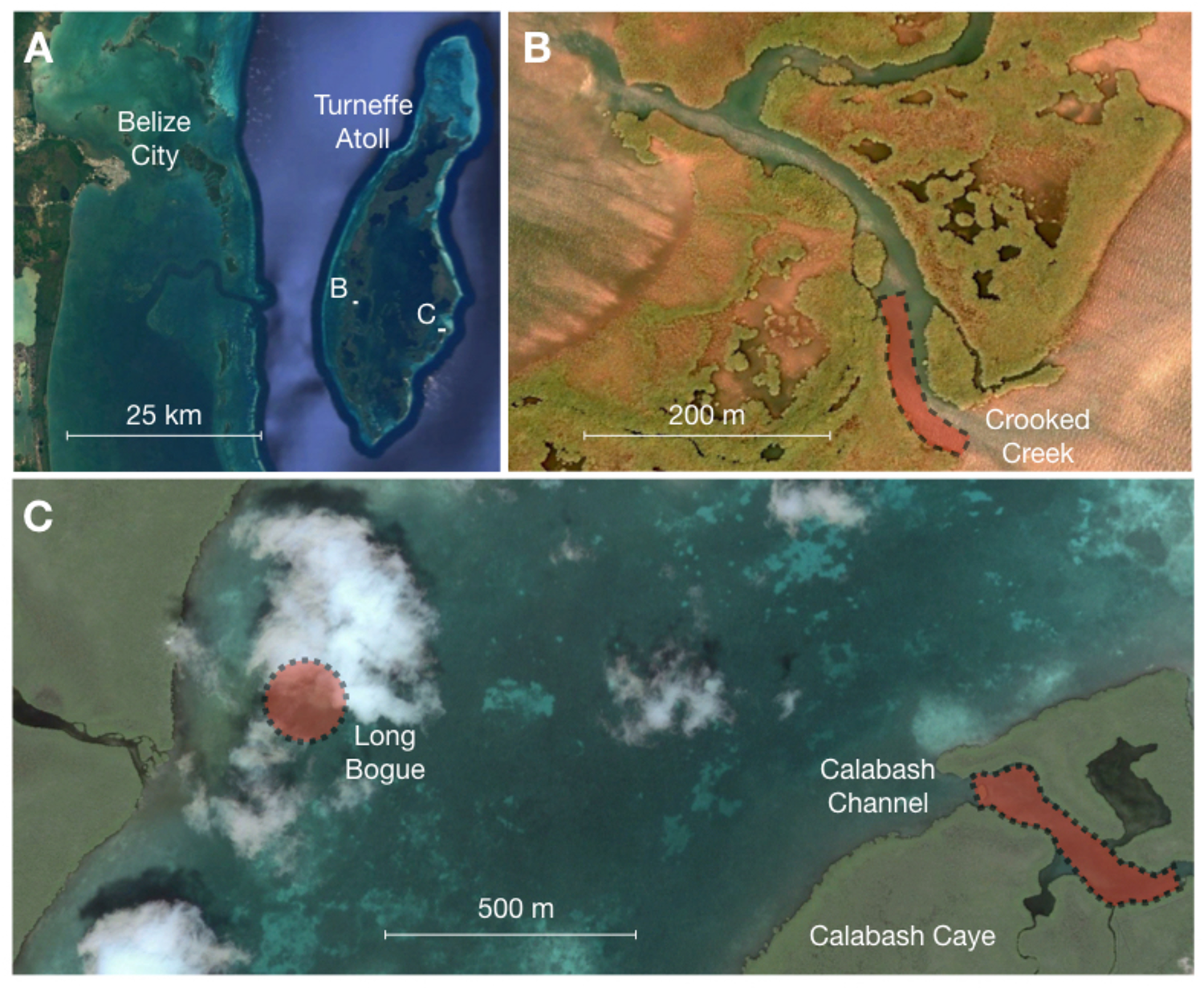
Location of study sites on Turneffe Atoll, Belize. (A) Turneffe Atoll. (B) Crooked Creek. Red shading indicates where corals were sampled growing on mangrove prop roots. The approximate center of the shoreline of Crooked Creek is 17°19’45.78”N and 87°55’6.35”W. (C) Lagoon patch reef site at Long Bogue and mangrove site at Calabash Channel. The approximate center of the Long Bogue site, where corals were sampled (red shading) is located at 17°17’25.84”N and 87°49’42.34”W. The approximate center of the Calabash Channel site, where corals were sampled growing on mangrove prop roots (red shading), is located at 17°17’16.23”N and 87°48’48.20”W. The image is dated August 26, 2005 and was obtained using Google Earth Pro (v 7.3.2.5776). Image © 2020 Maxar Technologies.

**Figure 2.**
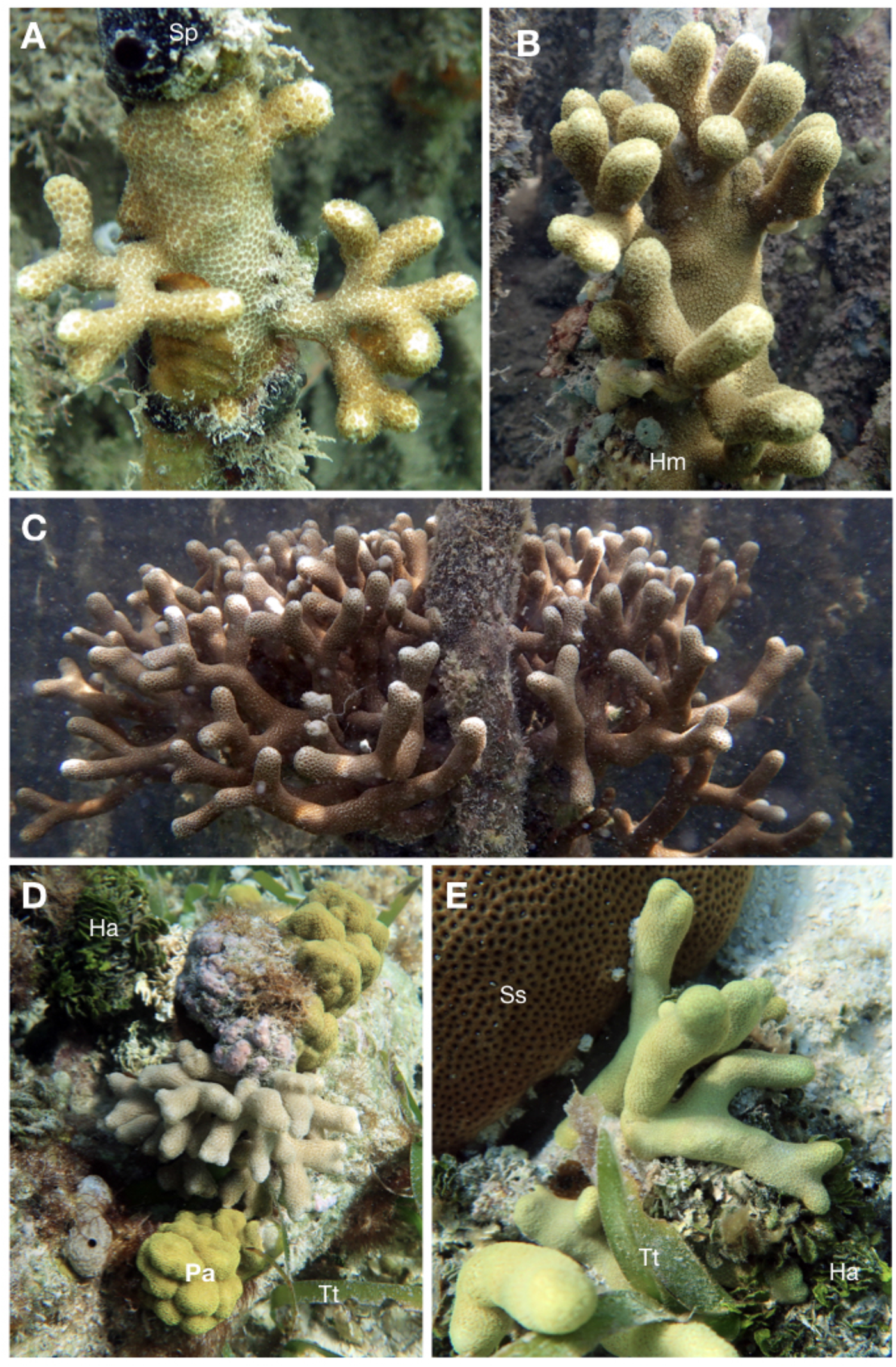
Representative photos of *Porites divaricata* from mangrove and lagoon habitats. (A-C) Corals growing on mangrove prop roots in Calabash Channel. (D-E) Corals growing in patch reef habitat in Long Bogue. Co-occurring organisms visible in these photos include: *Ha=Halimeda* sp.; *Hm=Haliclona manglaris; Pa=Porites astreoides; Ss=Siderastrea siderea;* Sp=*Spongia pertusa*; Tt=*Thalassia testudinum*. Photos by John R. Finnerty.

**Figure 3.**
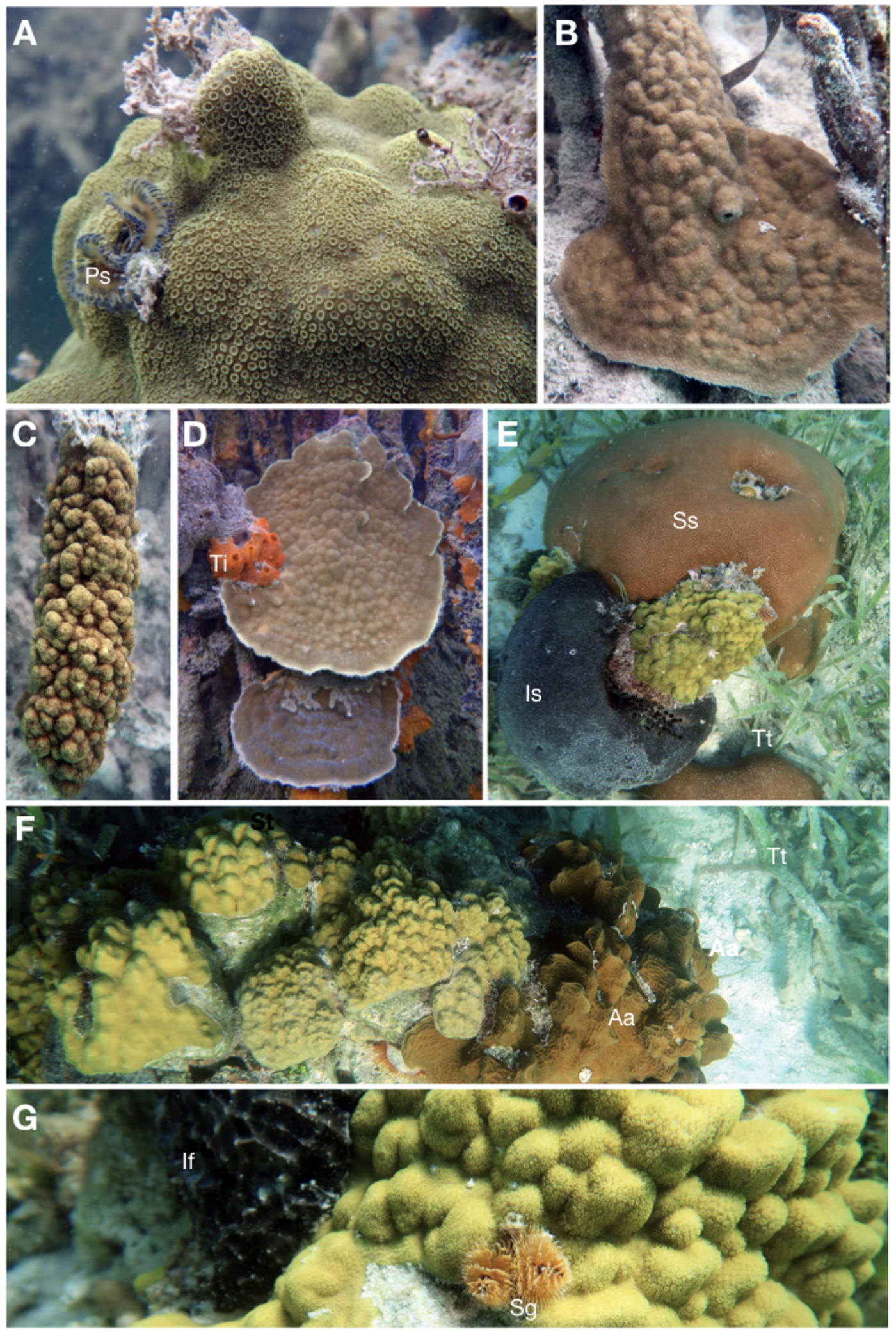
Representative photos of *Porites astreoides* from mangrove and lagoon habitats. (A) Coral growing on mangrove root in Calabash Channel. (B-D) Corals growing on mangrove roots in Crooked Creek. (E-G) Corals living in lagoon at Long Bogue. Co-occurring organisms visible in this photograph include: Aa=*Agaricia agaricites*; Ps=*Pomatostegus stellatus*; St=*Spongia tubulifera*; If=*Ircinia felix*; Is=*Ircinia strobalina*; Ss=*Siderastrea siderea*; Sp=*Spirobranchus giganteus*. Photos by John R. Finnerty.

While previous studies have explored the phenotypic variation of *P. astreoides* (Gleason 1993) and *P. divaricata* (Bengtsson et al. 2019) within habitat types, this study characterized differences between coral species inhabiting mangrove and lagoon habitats. Given that mangrove and lagoon habitats differ in key environmental parameters including temperature and light, we hypothesized that colonies from the two study locations would exhibit consistent and predictable differences in phenotype. Specifically, we expected that corals in the mangrove sites would exhibit traits associated with lower light levels, such as darker coloration (Gleason 1993) and distinct corallite architecture (Ow & Todd 2010, Soto et al. 2018). We also expected colony morphology to differ because in the mangroves, colonies are attached to an approximately vertical substrate (prop roots of the red mangrove, *Rhizophora mangle*), while in the lagoon colonies are attached to an approximately horizontal substrate. Over two years of sampling, we characterized multiple phenotypes of 249 coral colonies and found significant differences in colony size, structural complexity, color and form between corals inhabiting mangrove and lagoon habitats.

## 2. MATERIALS AND METHODS

### 2.1. Study site

Corals were characterized at three sites on Turneffe Atoll, Belize: a shallow lagoon site, and two mangrove-lined channels (Figure 1). Turneffe Atoll, located approximately 32 km off the coast of Belize, is one of only four atolls in the Western Hemisphere and is the largest of the three Belize atolls, with a surface area of approximately 531 km^2^ (Stoddart 1962). Unique among the Belizean atolls, Turneffe is dominated by mangroves. *Rhizophora mangle* is practically ubiquitous at the water’s edge, and lesser amounts of *Avicennia germinans* and *Laguncularia racemosa* grow in the interior of the hundreds of small mangrove islands known as “cayes.” The single lagoon site, hereafter called Turneffe Lagoon, consists of a shallow patch reef (0.5-3.0 m deep) located in Long Bogue, just west of Calabash Caye. One of the two mangrove sites, called Calabash Channel, is located ~1.4 km east of the Turneffe Lagoon patch reef. Within Calabash Channel, we assessed corals growing on the prop roots of *Rhizophora mangle* in three locations: (1) along the northern shore, (2) along the circumference of a small island, and (3) along the banks of a small creek that connects Calabash Channel to an expansive pond located within the interior of Calabash Caye. The second mangrove site we surveyed was at Crooked Creek, located on the western side of Turneffe Atoll, ~10.5 km west of the Turneffe Lagoon patch reef. Here, we monitored corals growing on *R. mangle* roots and/or on the mangrove peat bank along an approximately 150m stretch at the eastern edge of the creek where it enters the central lagoon of Turneffe Atoll (Figure 1b).

### 2.2. Environmental Monitoring

From November 2017 to November 2018, five Onset^®^ HOBO^®^ Pendant data loggers were deployed to record both temperature (°C) and light intensity (lux) every 3 hours. Three loggers were placed in Calabash Channel, dispersed among coral colonies that were surveyed. The loggers were affixed to bare mangrove roots (unshaded by epibionts) using nylon cable ties at approximately 40 cm depth, where mangrove corals are commonly found. Two loggers were placed in the Turneffe Lagoon patch reef site. For the light measurements, we utilized only the first seven days of recorded data as the instruments’ sensitivity to light declined rapidly due to fouling by marine organisms. Data collected between 2000 hrs and 0500 hrs were removed (all values were 0 lux during this time window). Lux was converted to photosynthetic photo flux density (PPFD: μmol m^-2^ s^-1^) with a standard sunlight conversion factor of 0.0185. We recorded the depth of each colony at each site in 2018. In the lagoon, colony depth was measured with a Laylin Speedtech SM-5 portable sounder and depth meter. In the mangroves, colony depth was measured from the water’s surface to the highest point of the colony using a measuring tape.

### 2.3. Linear Colony Dimensions and Ecological Volume

For each coral growing in the lagoon, height was measured as the vertical distance from the tallest part of the colony to its attachment with the substrate. Length and width were measured in a plane parallel to the substratum, with length representing the greatest linear dimension parallel to the substrate, and width measured at a point rotated 90° from where the length was taken. For corals growing on mangrove roots, linear dimensions were measured as previously described (Bengtsson et al. 2019, Scavo Lord et al. 2020). Height was measured as the perpendicular distance from the root to the most distant extent of the colony. Length was measured as the linear extent of the colony along the root, and width was measured at 90° to both height and length. From these linear dimensions, we calculated ecological volume as πHr^2^ where H is equal to colony height, and r is equal to (colony width + colony length)/4 (Shaish et al. 2006).

### 2.4. Colony Color

Corals were photographed *in situ* against a laminated DKK color standard (www.dgkcolortools.com) that included a six-step gray scale featuring true white, as well as 12% and 18% gray. Using the method presented by Winters et al. (2009), these photographs were used to estimate the chlorophyll density of each coral colony. Briefly, all photographs were standardized using the gray scale and the MATLAB macro “CalibrateImageA.” Then, the MATLAB macro “AnalyzeIntensity” was used to calculate mean red channel intensity for 20 swatches of 25×25 pixels for each coral. Higher red channel intensity values indicate fewer algal photosynthetic pigments (Winters et al. 2009).

### 2.5. Branch Number, Spacing, and Thickness in P. divaricata

Branch thickness and branch number were recorded in the field while snorkeling. Branch thickness was measured using calipers at a point 2.5 cm below the branch tip for three random branches from each colony, and the average value was calculated. For branch number, only visibly healthy branches at least ~1 cm in length were counted. Counts were performed a minimum of two times for each coral colony, and the count was repeated if there was a discrepancy. Branch spacing (branch tips cm^-3^) was calculated by dividing branch number by ecological volume.

### 2.6. Rugosity and Form in P. astreoides

Surface rugosity of *P. astreoides* was determined using the bar and chain method (Risk 1972) at each colony’s widest point. Based on *in situ* observations corroborated by photographs, the form of each colony was characterized as mounding, plating, or a combination of mounding and plating.

### 2.7. Corallite Density and Area

For both species, a flexible stencil of known area was placed flush against each coral’s surface to obtain three macrophotographs of the corallites circumscribed by the stencil. In *P. divaricata*, the three photographs were taken at a point 1.0 cm below the branch tip on three random branches. In *P. astreiodes*, the photographs were taken at three random locations on the colony. From the photographs, corallite density and corallite area were determined. Corallite density was measured by counting all corallites whose central point was located within the circular frame and dividing by the area of the frame. The multipoint tracking tool of ImageJ (version 1.5a) was used to count the corallites present in the frame (Abramoff et al. 2004). For each coral, the average corallite density was determined from the three photos taken. Corallite area was determined by manually tracing corallites in the photographs and calculating the area of the irregular shape using ImageJ. Only corallites located entirely within the circular frame were included in the analysis. For each coral, the average corallite area was determined for the three photos taken.

### 2.8. Statistical Analysis of Coral Phenotypes

For all variables, we compared mean values for mangrove versus lagoon samples from each species. Data from Calabash Channel and Crooked Creek were combined to generate an average value for mangrove specimens. If the raw data or logarithmically transformed data met the assumption of normality according to a Shapiro-Wilkes test, an independent-sample *t*-test was conducted to compare means. If the data violated the assumption of normality even following ln transformation, a Mann-Whitney-Wilcoxon test (non-parametric) was used to compare means. Each test was conducted both with outliers and without outliers (if present), which were removed using the *boxplot$out* command in R (R Core Team 2018), which identifies the values lying further than 1.5 times the interquartile range from the upper and lower quartiles.

## 3. RESULTS

### 3.1. Number of Colonies Assessed by Habitat, Site, Species and Year

Over the course of two sampling periods (November 2018, 2019), we obtained data from 249 distinct coral individuals: 155 specimens of *P. divaricata* and 94 specimens of *P. astreoides* (Table 1). The 146 specimens assessed in 2018 were used for the analyses of ecological volume, branch number, branch spacing, branch thickness, and rugosity. The 103 specimens assessed in 2019 were used for the analyses of corallite area, corallite density, and colony color.

**Table 1.**
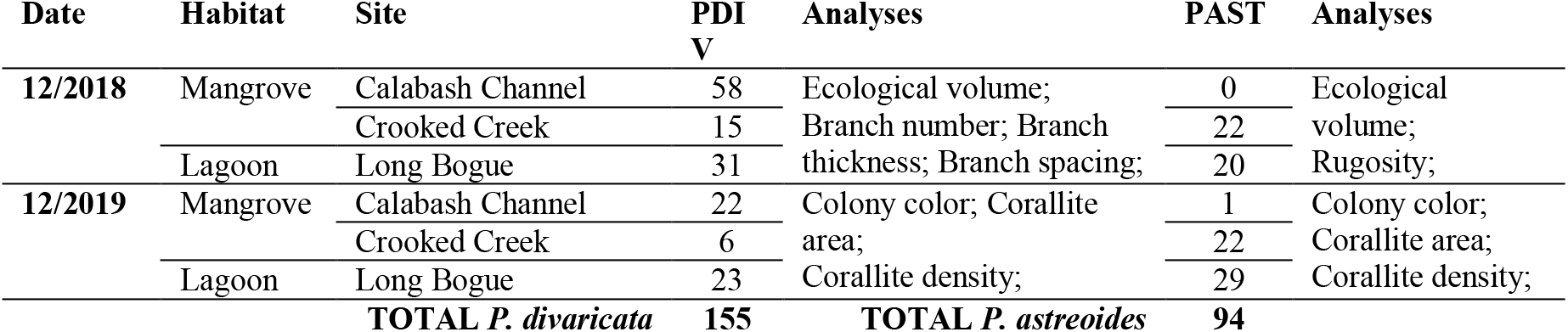
Specimens analyzed by habitat, site and species.

| Date | Habitat | Site | PDI<br>V | Analyses | PAST | Analyses |
| --- | --- | --- | --- | --- | --- | --- |
| 12/2018 | Mangrove | Calabash Channel | 58 | Ecological volume; | 0 | Ecological |
|  |  | Crooked Creek | 15 | Branch number; Branch | 22 | volume; |
|  | Lagoon | Long Bogue | 31 | thickness; Branch spacing; | 20 | Rugosity; |
| 12/2019 | Mangrove | Calabash Channel | 22 | Colony color; Corallite | 1 | Colony color; |
|  |  | Crooked Creek | 6 | area; | 22 | Corallite area; |
|  | Lagoon | Long Bogue | 23 | Corallite density; | 29 | Corallite density; |
| TOTAL *P. divaricata* |  |  | 155 | TOTAL *P. astreoides* | 94 |  |

### 3.2. Environmental Variation Across Habitats

The mangrove and lagoon habitats were significantly different with respect to temperature, light, and the average depth at which corals were located (Figure 4). From November 2017 to December 2018, the average temperature in the mangroves of Calabash Channel was significantly warmer than in the lagoon at Long Bogue (Figure 4A; 28.77 ± 1.84 °C vs. 28.62 ± 1.79 °C (mean ± SD); p=0.038, respectively). Furthermore, the average daily temperature variance at Calabash Channel was significantly greater (Figure 4B; 0.58 ± 0.39 °C vs. 0.24 ± 0.17 °C, respectively; p<0.001). The average light levels were significantly greater in the lagoon than in the mangroves (Figure 4C; 165.39 ± 64.45 μmol m^-2^ s^-1^ vs. 54.50 ± 47.96 μmol m^-2^ s^-1^, p<0.001), as was the daily variation in light levels (Figure 4D; 38,085.62 ± 41,957.42 vs. 8,180.02 ± 13,606.61; p<0.001). For both species, mean colony depth was significantly greater in the lagoon than in the mangroves (Supplemental File 1).

**Figure 4.**
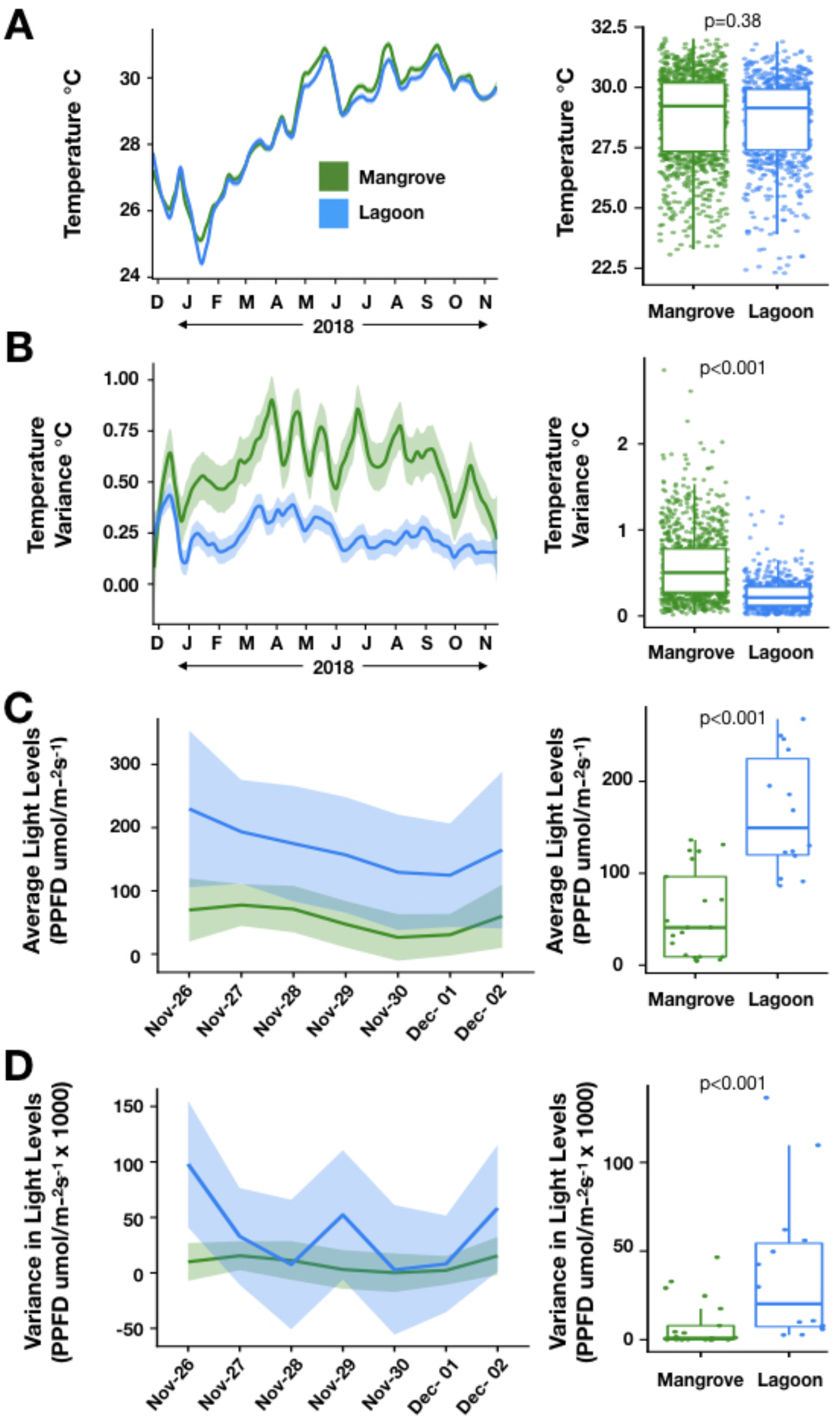
Environmental differences between lagoon and mangrove sites. A) Mean daily temperature tracked over 12-month period (left) and averaged over the entire period (right). (B) Daily temperature variance tracked over a 12-month period (left) and averaged over the entire period (right). (C) Mean daily light levels tracked over a 7-day period (left) and averaged over the entire period (right). (D) Daily light variance tracked over a 7-day period (left) and averaged over the entire period (right). In the panels on the left, the shaded areas around the lines indicate standard error. In the panels on the right, the line dividing the line dividing the box represents the median, and the upper and lower sides of the box denote the first and third quartiles. The whiskers extend up to 1.5 times the interquartile range of the box, and dots past the whiskers are outliers.

### 3.3. Colony Size

For both coral species, the mean ecological volume was greater for the lagoon samples than the mangrove samples (Figure 5), though the difference was statistically significant only for *P. divaricata* (p<0.001, regardless of whether we included or excluded outliers; compare Figure 5A vs 5B). The mangrove samples for *P. divaricata* exhibited by far the greatest size range, from 3.62 to 18117.25 cm^3^. *P. astreoides* from the lagoon exhibited the next greatest size range, from 271.89 to 16679.96 cm^3^ (Figure 5C).

**Figure 5.**
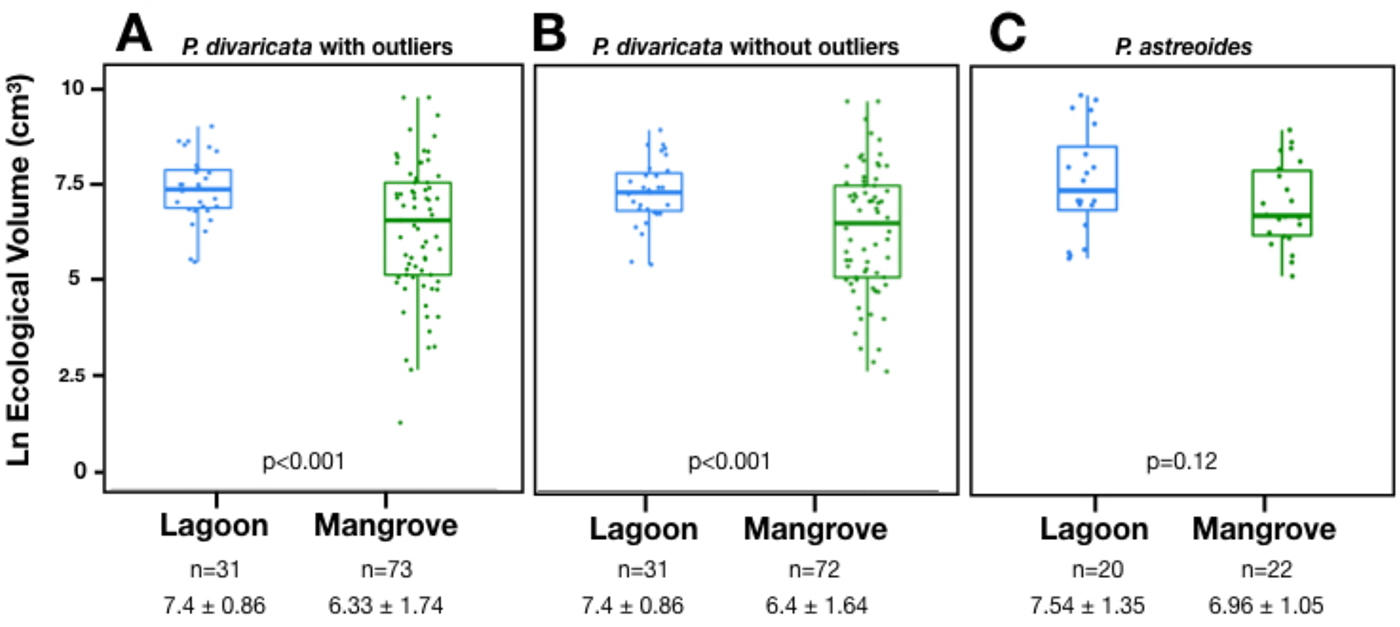
Ecological volume of *P. divaricata* and *P. astreoides* in lagoon and mangrove environments. Each point on the graph represents one colony. The line dividing the box into two sections represents the mean, the top and bottom lines of the box represent the interquartile range, and the whiskers represent the largest and smallest value that lie within 1.5 times the interquartile range. Box plots (A) and (B) show data for *P. divaricata* with (A) including outliers while (B) does not. Panel (C) shows the data for *P. astreoides*. Outliers were tested for in *P. astreoides*, but none were detected. For both species, statistically significant differences in ecological volume between sites were determined with an independent two-sample *t*-test.

### 3.4. Colony Color Intensity

For both coral species, the mangrove colonies exhibited significantly greater intensity in the red color channel than lagoon colonies (Figure 6). The differences between mangrove and lagoon colonies were statistically significant at p<0.001 for both *P. divaricata* (Figure 6A,B) and *P. astreoides* (Figure 6C), regardless of whether statistical outliers were included (Figure 6A,C) or excluded (Figure 6B). In *P. divaricata*, the range of color intensity values was greater in the mangroves than in the lagoon (54.5-173.6 vs. 90.3-190.5). In *P. astreoides*, the range of color intensity values was greater in the lagoon (93.6-179.5 vs. 46.1-113.9). For these analyses. the intensity for the red channel was used because it is known to have the highest correlation with chlorophyll density (Winters et al. 2009); however, mangrove corals also exhibited significantly greater color intensity in both the green and blue channels (Supplemental File 2).

**Figure 6.**
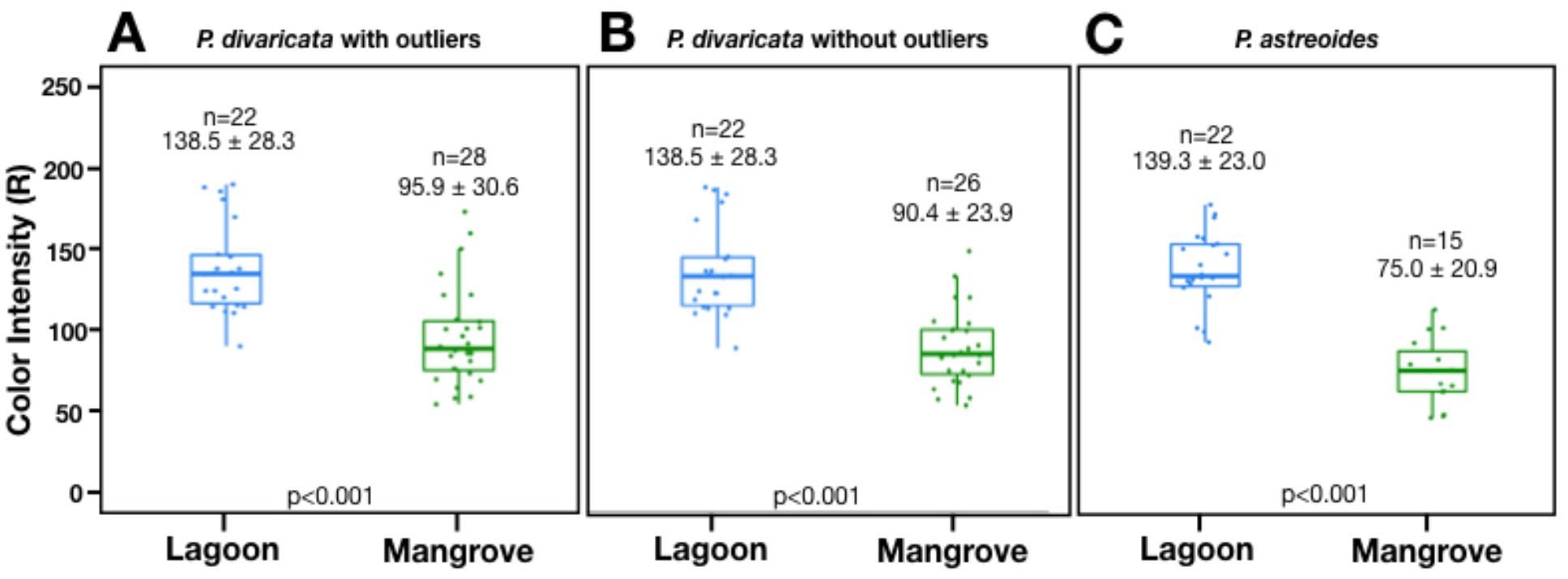
Colony color of *P. divaricata* and *P. astreoides* in lagoon and mangrove environments. Red channel intensity is a proxy for chlorophyll pigment density, with higher intensity values corresponding to lower chlorophyll densities. Mean red channel intensity in *P. divaricata*, including (A) and excluding (B) outliers. Mean red channel intensity in *P. astreoides* (C) in which no outliers were detected. The data are represented as in Figure 5. Statistically significant differences in colony color between sites were determined with a Mann-Whitney-Wilcoxon test in *P. divaricata*, and an independent two-sample *t*-test in *P. astreoides*. An independent *t*-test was also performed on ln transformed *P. divaricata* values, because transformation resulted in normalized data (p<0.001); however, in order to display the values on the same scale as *P. astreoides*, we conducted the analysis on the non-transformed data.

### 3.5. Branch Metrics

Branch number was the only branch metric that exhibited a clear difference between mangrove and lagoon *P. divaricata* (Figure 7). On average, lagoon colonies had about twice as many branches as mangrove colonies (Figure 7C; 44.8 versus 22.2 branches), and more than three times as many if we removed the outliers from the mangrove sample (Figure 7D). These differences were highly significant, whether outliers were included or excluded (p<0.001). Mangrove colonies exhibited a greater range of branch numbers (1-157) than lagoon colonies (10-83).

**Figure 7.**
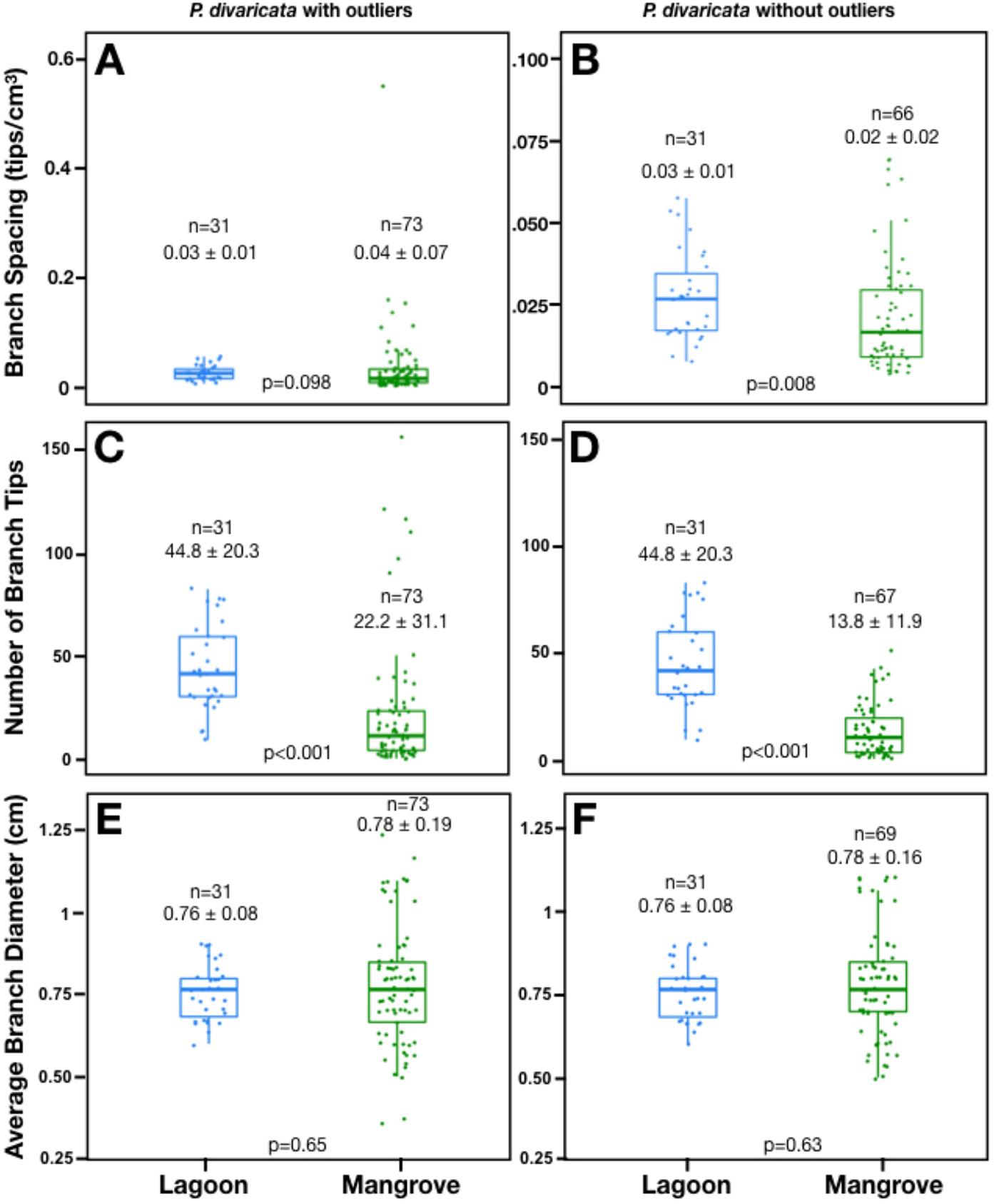
Branch number, spacing, and thickness in *P. divaricata*. Mean branch number, including (A) and excluding (B) outliers. Mean branch spacing, including (C) and excluding (D) outliers. Mean branch thickness, including (E) and excluding (F) outliers. The data is represented as in Figure 7. Statistically significant differences in all three branch-metrics between sites was determined with a Mann-Whitney-Wilcoxon test.

In contrast to branch number, there was no significant difference in average branch thickness between mangrove and lagoon habitats, with or without outliers (Figure 7E,F). However, we again observed a greater range of values in the mangroves (0.35-1.2 cm) than in the lagoon habitat (0.6-0.9 cm).

Branch density did not differ between mangrove and lagoon samples unless outliers were removed (Figure 7A,B). With outliers included, the mean branch density of lagoon corals was less than that of mangrove corals (0.03±0.01 vs. 0.04±0.07 branches cm^-3^; p=0.12; Figure 7A). With outliers excluded, the mean branch density of lagoon corals exceeded that of mangrove corals (0.03 ± 0.01 vs. 0.02 ± 0.02 branches cm^-3^, p=0.008; Figure 7B). Mangrove colonies showed a higher range in branch density values than lagoon colonies (0.004-0.55 branches cm^-3^ versus 0.01-0.06 branches cm^-3^ respectively).

### 3.6. Rugosity and Form

*P. astreoides* exhibited significantly higher mean rugosity in the lagoon than in the mangrove, regardless of whether outliers were included (p=0.04; Figure 8A) or excluded (p=0.02; Figure 8B). When outliers were included, the range of rugosity was greater among mangrove corals (1.07-2.46) than lagoon corals (1.04-2.09). The lagoon site exhibited much greater consistency in form, with 100% of colonies surveyed exhibiting a mounding phenotype (Figure 8C). The majority of mangrove corals (59%) exhibited a plating morphology (e.g., Figure 3D), while only 27% exhibited a mounding form (e.g., Figure 3C), and 14% exhibited a combination of plating and mounding (e.g., Figure 3B).

**Figure 8.**
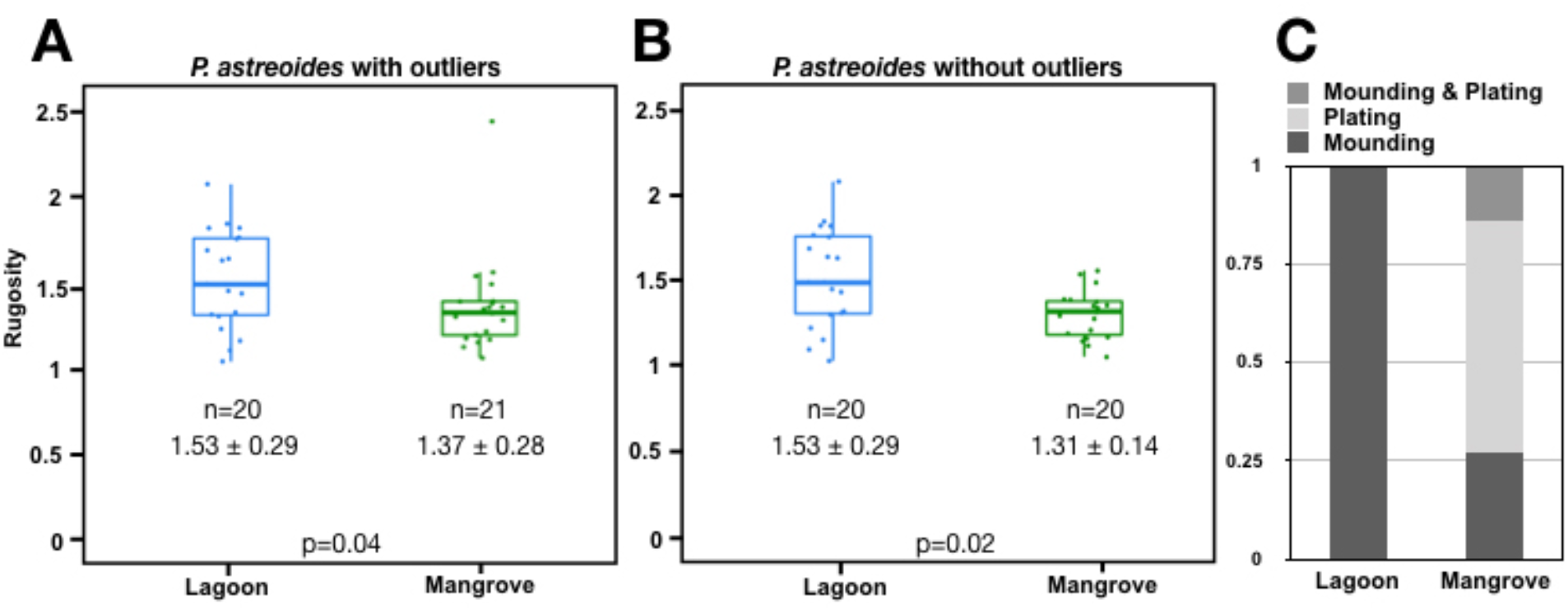
Rugosity and form in *P. astreoides*. Mean rugosity, including (A) and excluding (B) outliers. (C) Relative abundance of three *P. astreoides* colony forms: (plating, encrusting, or a combination of both. Examples of each colony form: (D) a mounding colony on a mangrove root; (E) two plating colonies on a mangrove root; (F) a mounding colony in a lagoon habitat; (G) a colony on a mangrove root displaying a combination of mounding and plating forms. Statistically significant differences in rugosity between sites were determined with a Mann-Whitney-Wilcoxon test.

### 3.7. Corallite Morphology

In both species, average corallite density was significantly greater in the mangrove sample than the lagoon sample (p<0.001), regardless of whether outliers were included (Figure 9A,E) or excluded (Figure 9B,F). For *P. divaricata*, mangrove colonies exhibited a greater range of corallite density values (57.7-138.0 corallites cm^-2^) than lagoon colonies (46.6-109.4 corallites cm^-2^; Figure 9A,B). The same was true for *P. astreoides*, with ranges of 51.4-118.2 and 46.0-75.8 corallites cm^-2^ in the mangrove and lagoon, respectively (Figure 9E,F).

**Figure 9.**
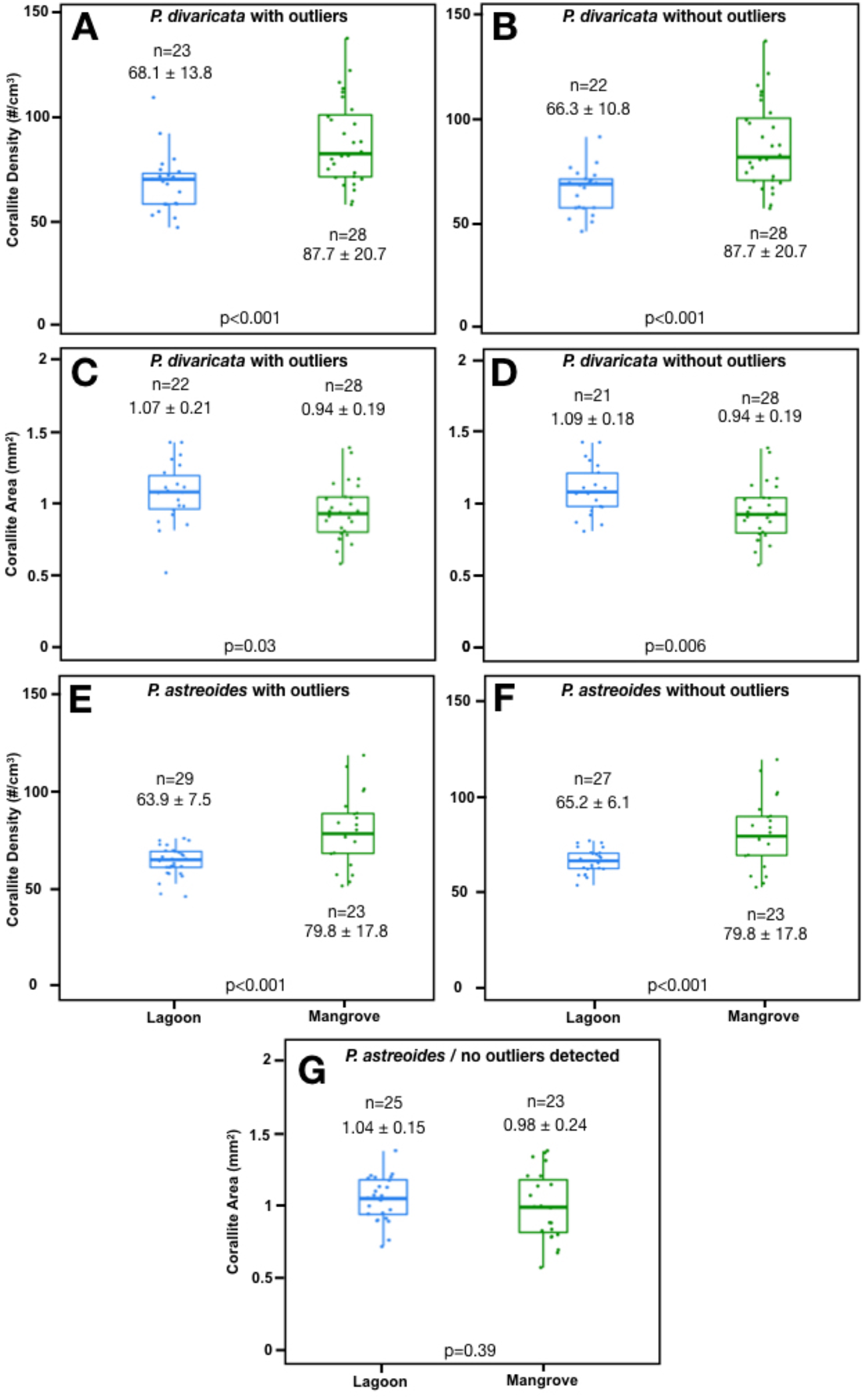
Corallite area and density in *P. divaricata* and *P. astreoides* from lagoon and mangrove environments. Mean corallite density in *P. divaricata*, including (A) and excluding (B) outliers. Mean corallite area in *P. divaricata*, including (C) and excluding (D) outliers. Mean corallite density in *P. astreoides*, including (E) and excluding (F) outliers. Mean corallite area in *P. astreoides* (G) for which no outliers were detected. The data is represented as in Figure 7. Statistically significant differences in corallite density between sites for *P. divaricata* was determined with a Mann-Whitney-Wilcoxon test; although, after outliers were removed, the data were normally distributed and independent two-sample was also performed, producing the same results. Statistically significant differences in corallite density in *P. astreoides*, and corallite area in both *P. divaricata* and *P, astreoides* were determined with an independent two-sample *t*-test.

Average corallite area was slightly higher in lagoon versus mangrove samples of both corals (Figure 9C,D,G), although only the difference in *P. divaricata* was statistically significant (p=0.03/0.006, including or excluding outliers). The range of corallite areas for the lagoon samples of *P. divaricata* (0.52-1.42) was slightly higher than the corresponding values for the mangrove samples (0.58-1.38). while the opposite trend was seen in *P. astreoides* (0.56-1.37 and 0.71-1.37 for the mangroves and lagoon, respectively).

## 4. DISCUSSION

Mangroves are increasingly being recognized as an important component of the ecological niche for many coral species (Yates et al. 2014, Hernández Fernández 2015, Rogers 2017, Bengtsson et al. 2019, Camp et al. 2019, Scavo Lord et al. 2020). Here, we demonstrate significant differences in key environmental variables between the mangrove and adjacent coral-supporting habitats. This suggests that coral species exploiting this range of habitats may be capable of expressing a wider range of phenotypes than those which cannot. Indeed, we document a number of statistically significant differences in the phenotype of coral colonies in mangrove and lagoon habitats, including differences in colony size, color, structural complexity, and corallite morphology. For each phenotypic trait, the direction of the difference between mangrove and lagoon corals was consistent in *P. divaricata* and *P. astreoides*.

With respect to size, lagoon corals exhibited a larger mean ecological volume than mangrove corals, though the difference was statistically significant only in *P. divaricata*. Differences in overall colony size could reflect size constraints on corals growing in the mangroves. In the mangrove habitats studied, colonies were found growing primarily on prop roots, which can be either grounded in the peat bank or aerially suspended. On grounded roots, lateral expansion of colonies is typically constrained by other roots and/or root epibionts, and vertical expansion is constrained by the water’s surface and shallow peat bank. Aerial roots can extend outward from the peat bank, providing colonies more space to grow, and the largest colonies we observed were often situated on such roots. However, such large colonies are more directly exposed to the currents, putting the corals at greater risk of being dislodged during a storm. The colonies may also grow too heavy for the root to support. This could result in wholesale loss of the colony and possibly accelerated breakage of the root. This phenomenon has previously been observed during longitudinal monitoring of mangrove sponges (Bingham & Young 2005) and mangrove corals (Scavo Lord et al. 2020).

The lagoon might also support larger colonies as a result of greater energetic resources available via photosynthetic by-products of Symbiodiniaceae, as light availability has been correlated with increased growth rates in a number of coral species including *P. astreoides* (Huston 1985). We suspect that variability in light levels could explain the greater range of colony sizes in mangrove *P. divaricata*, but our existing light data lack the spatial resolution necessary to test this hypothesis. In the current study, three light meters were deployed adjacent to corals in typical mangrove locations, where they were shaded by the canopy throughout much of the day. As a result, the measured light levels were relatively low and invariant. While these shaded locations are where we most often observed corals, we have observed that the largest colonies tend to be found on roots that extend out from beneath the canopy, where they would be exposed to more light. Future studies could test this hypothesis by deploying a larger number of light meters immediately adjacent to mangrove corals spanning a wide range of sizes.

For both species, colonies in the lagoon exhibited greater color intensity than colonies in the mangroves. As higher intensity of the red channel is inversely correlated to chlorophyll density (Winters et al. 2009), this suggests mangrove colonies exhibit higher chlorophyll pigment density than conspecifics in the lagoon. Elevated chlorophyll concentration is a well-characterized adaptive response to maximize light harvesting by photosynthetic symbionts in corals inhabiting low-light environments (Abramovitch-Gottlib et al. 2005, Stambler & Dubinsky 2005). The same trend was documented in a comparison of mesophotic (45-50m) versus shallow (20-25m) colonies of *Montastrea cavernosa*, where mesophotic colonies contained significantly more Symbiodiniaceae cells, chlorophyll a per Symbiodiniaceae cell, and chlorophyll a and c2 per unit area (Polinski & Voss 2018). Similarly, increasing chlorophyll concentrations were observed in branch fragments of *Stylophora pistillata* transplanted to lower light levels (from 95% to 0.8% photosynthetic active radiation PAR0; Titlyanov et al. 2001).

The observed differences in colony form between habitats might also be attributable to variation in light (Todd 2008). In *P. astreoides*, the hemispherical mounding phenotype was found exclusively in the lagoon, while the plating, more flattened form was common in the mangroves. Variation in growth forms in differing light regimes is thought to be an adaptive response to maximize light capture in low light environments or enable self-shading in extremely high-light environments (Klaus et al. 2007). For example, colonies of *Porites rus* (formerly *Synaraea convexa*) exposed to high light levels exhibited hemispherical forms with short branches, while colonies exposed to the lowest light levels exhibited explanate or plating forms (Jaubert 1977). Similarly, the boulder star coral, *Orbicella annularis*, maximizes light capture in low light by growing in a flattened growth form (Dustan 1975).

Differences in corallite morphology were also consistent across habitats. Mean corallite density was higher in mangrove versus lagoon colonies for both species. Simultaneously, mean corallite area was also smaller in mangrove versus lagoon colonies, but this result was only significant in *P. divaricata*. Light availability has been linked to various aspects of corallite morphology. In particular, lower light is associated with decreasing corallite size (Beltran-Torres & Carricart-Gavinet 1993), as in *Montastrea cavernosa* where mesophotic colonies were found to have smaller corallites than shallow conspecifics (Studivan et al. 2019). Light availability can also influence the spacing between corallites, resulting in a greater degree of spacing between corallites in corals exposed to lower light (Studivan et al. 2019). While corallite spacing was not measured here, corallite density was significantly higher in mangrove representatives of both species. If corallite area stays the same between habitats, which it did in *P. astreoides*, this would suggest a reduction in spacing between corallites in mangrove corals, at odds with findings from other corals (Beltran-Torres & Carricart-Gavinet 1993). However, mangrove representatives of P. divaricata did exhibit significantly smaller corallite area, and smaller corallites or smaller polyps generally is suspected to maximize surface area for food capture (Sebens 1997). Smaller corallite size, and a higher density of corallites per unit area, might therefore suggest a greater reliance on heterotrophy in lower light environments, such as the mangroves. This reliance on heterotrophy may be particularly advantageous in nutrient-rich mangrove habitats, which could compensate for lower and more variable light availability. In the temperate coral *Cladocera caespitosa*, corals maintained in a low light environment used nutrients from heterotrophy to supplement calcification, and corals under high light converted carbon from feeding to tissue biomass (Hoogenboom et al. 2008). Similar heterotrophic plasticity was observed in a laboratory study of *Goniastrea retiformis*, which increased its feeding rate to fully compensate for reduced phototrophy when light levels were attenuated by suspended particulate matter; interestingly, under the same conditions, *Porites cylindrica* increased heterotrophy only slightly, and lost energy reserves as a result (Anthony & Fabricius 2000). Plasticity in trophic strategies could be an important survival strategy in corals that are able to exploit mangrove habitats, but the relative reliance on heterotrophy and autotrophy could impact different biological processes differently, as in the facultatively symbiotic *Astrangia poculata* where wound healing and total tissue cover were impacted differently by autotrophy and heterotrophy (Burmester et al. 2018).

Phenotypic variability has been widely documented in corals spanning a range of environmental gradients. The consistent phenotypic differences across habitat types in the two species described here could be driven by constraints imposed by the habitat, phenotypically plastic responses to environmental variation, genetic differences due to the selection of locally advantageous phenotypes or by any combination of the above. While more research is needed to determine the degree to which phenotypic plasticity is driving the observed differences (i.e. reciprocal transplants of coral genets between habitats), we suspect that it plays a prominent role in the variation described here. First, all phenotypic differences observed between sites are consistent with documented trends in other coral species in low vs. high light environments.

Importantly, there are many other environmental parameters that are known to drive phenotypic variation, including flow (Kaandorp et al. 1996, Bruno & Edmunds 1998), sedimentation (Todd et al. 2001), and nutrient levels (Bongiorni et al. 2003a, Bongiorni et al. 2003b). These other factors are also likely to contribute to variation in phenotype between mangrove and non-mangrove corals of the same species, as mangroves are generally more stagnant, turbid, and nutrient-rich compared to reefs (Nagelkerken et al. 2008, Granek et al. 2009). Additionally, with nearly every phenotypic trait, mangrove corals exhibited greater variability. This may reflect greater micro-environmental variation in the mangroves than the lagoon. For example, different prop roots can be exposed to very different light levels or flow patterns depending on their distance from the peat bank, canopy, or other roots (Farnsworth & Ellison 1996).

Corals that are “less solely dependent on reef habitats” have been shown to be less vulnerable to environmental change (Carpenter, 2008). This may be attributed to (1) their capacity to tolerate a greater range of environmental variation within reef habitats, and (2) their ability to exploit non-reef habitats when conditions on the reef deteriorate. One manifestation of wider environmental tolerances is greater phenotypic plasticity. Weedy corals have been found to exhibit greater intraspecific variation in phenotypic traits than corals with other life-history strategies, such as competitively-dominant species, stress-tolerant species, and generalists (Darling et al. 2012). The fact that “weedy” corals are increasing in relative abundance on reefs could be due, in large part, to this intraspecific variation; however, there could be several other contributors to their success, such as a brooding reproductive mode and high fecundity (Darling et al. 2012).

This is the first study to directly compare the morphology of corals across mangrove and shallow patch-reef habitats. Given increasing interest in the role of non-reef habitats for supporting coral resilience (Yates et al. 2014, Hernández Fernández 2015, Rogers 2017, Bengtsson et al. 2019, Camp et al. 2019, Scavo Lord et al. 2020), it is important to understand whether and how coral phenotypes can change to accommodate the conditions found in what have historically been regarded as suboptimal habitats for corals, such as mangroves. Going forward, it will be important to differentiate genetic from environmental factors in the phenotypic diversity documented here. Towards this end, ongoing research is investigating (1) the effects of transplanting corals within and between mangrove and lagoon habitats and (2) the genetic differences between mangrove and lagoonal populations of *Porites* collected from the sites used in this study (Scavo Lord et al. in preparation).

## ACKNOWLEDGEMENTS

This research described here was supported by National Science Foundation grants DGE-1247312 to KSL and IOS-1354935 to JRF. Fieldwork conducted in 2018 was supported by Boston University through the Graduate Research Abroad Fellowship to KSL. The research was conducted under Aquatic Scientific Research Permits #0041-18 (2018), and #0064-19 (2019) issued by the Belize Fisheries Department. We would like to thank Felicia Cruz and Mauro Gongora at the Belize Fisheries Department for assistance in obtaining these research permits. We would also like to thank Julia Hammer-Mendez, Jonathan Perry, and Justin Scace, staff of the Boston University Marine Program, for their logistical and technical support in the Marine Semester. We are grateful for the expert technical assistance and natural history knowledge of boat captains, staff, and researchers at Calabash Caye Field Station and the University of Belize (including Berris Torres, Derron Torres, Joshua Morey, and Javon Castillo). We are also grateful to the Bertarelli Foundation and the Oak Foundation for their contributions to the research infrastructure at Calabash Caye Field Station, which facilitated the marine conservation related research described here.

---

**Supplementary File 1.**
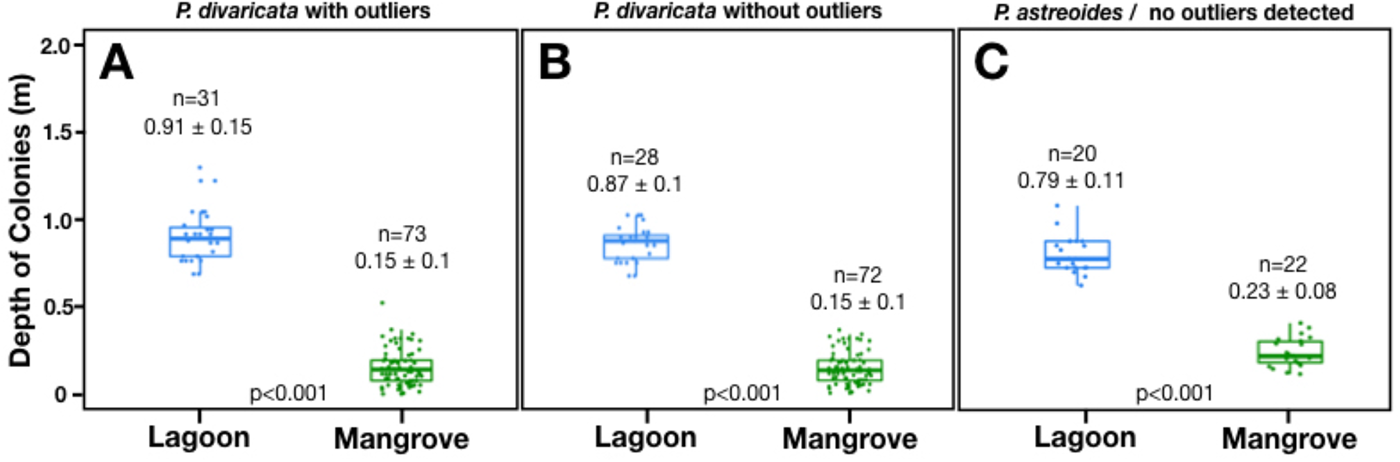
Colony depth between sites. Mean colony depth in *P. divaricata*, including (A) and excluding (B) outliers. Mean colony depth in *P. astreoides* (C) in which no outliers were detected. The data is represented as in Figure 7. Statistically significant differences in colony depth between sites was determined with a Mann-Whitney-Wilcoxon test in *P. divaricata*, and an independent two-sample *t*-test in *P. astreoides*.

**Supplementary File 2.**
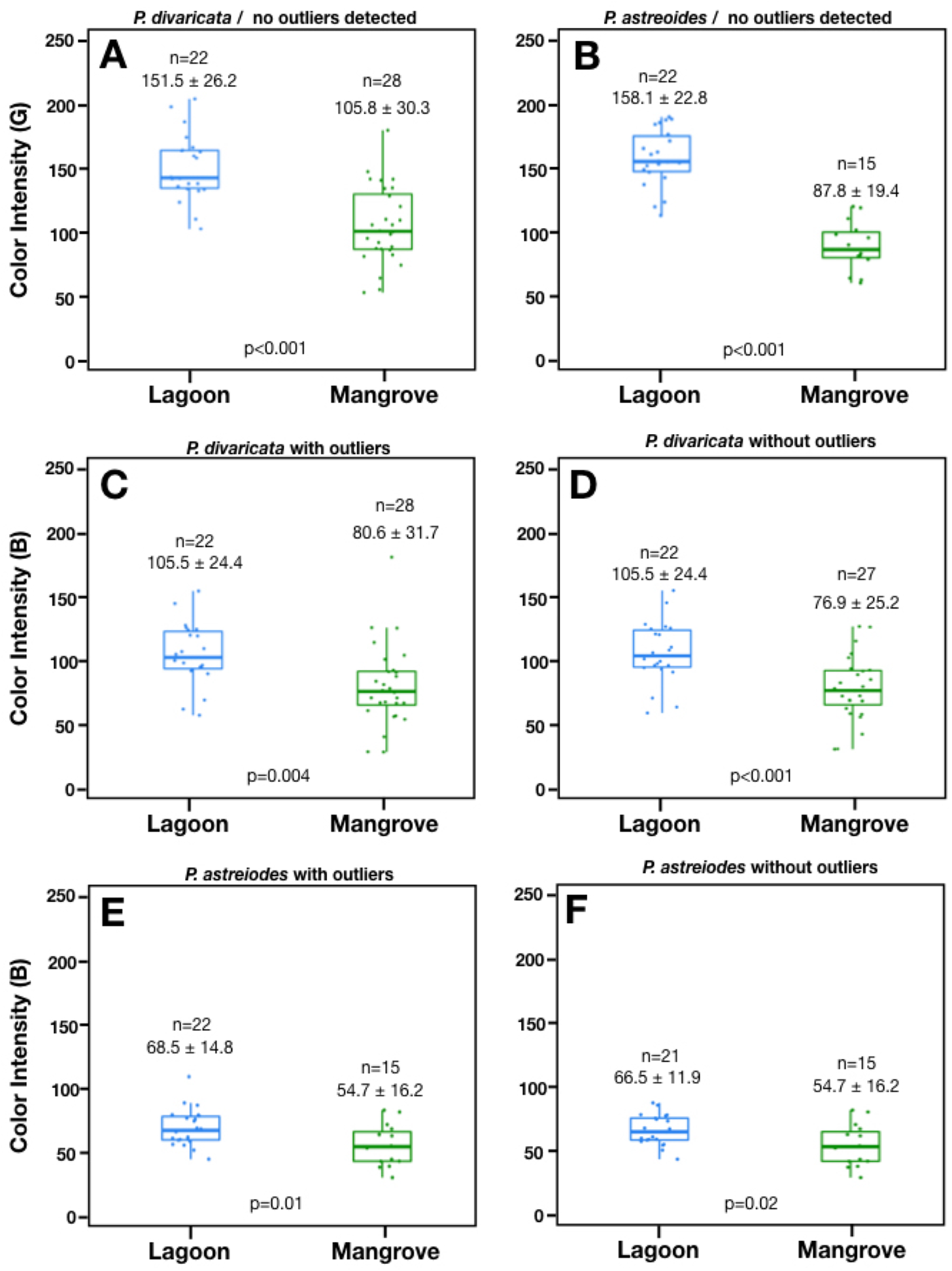
Color intensity of the green and blue channels in *P. divaricata* and *P. astreoides*. Mean color intensity in the green (G) color channel in *P. divaricata* (A) and *P. astreoides* (B) in which no outliers were detected. Mean color intensity in the blue (B) color channel in *P. divaricata* with (C) and without (D) outliers, and *P. astreoides* with (E) and without (F) outliers. The data are represented as in Figure 5. Statistically significant differences in colony color intensity in the green and blue channels between sites was determined with an independent two-sample *t*-test in both species.

